# Maternal E-Cigarette Exposure Exacerbates Neonatal Ischemic Brain Injury Through the miR-30c-5p–Neurod1 Axis

**DOI:** 10.64898/2026.11.09.765511

**Authors:** Yanyan Zhang, Rongjun Yu, Jia Tian, Wanshu Yu, Siyi Jiang, Jiazichao Tu, Qingyi Ma, Ruize Zhang, Shuwei Sun, Lubo Zhang, Daliao Xiao, Yong Li

## Abstract

**Background:** Neonatal ischemic stroke is an important cause of mortality and long-term neurological morbidity in infants. Maternal smoking and e-cigarette exposure are recognized risk factors for adverse neonatal outcomes, but the mechanisms underlying increased susceptibility to ischemic brain injury remain unclear. This study investigated whether dysregulation of the miR-30c-5p–Neurod1 axis contributes to this susceptibility.

**Methods:** Pregnant rats were exposed to e-cigarette aerosols from embryonic day 4 to 20. Studies were conducted in male offspring on postnatal day 10. Baseline molecular changes were assessed using microRNA sequencing, quantitative reverse-transcription polymerase chain reaction, Western blotting, RNA-induced silencing complex (RISC) and luciferase reporter assays. Ischemic injury was evaluated following transient middle cerebral artery occlusion (tMCAO). Flow cytometry and molecular analyses were performed to characterize associated cellular changes. The effects of in vivo miR-30c-5p mimic administration, Neurod1 knockdown, and miR-30c-5p inhibition were also investigated.

**Results:** Maternal e-cigarette exposure increased tMCAO-induced infarct size and was associated with increased apoptosis and senescence-associated changes, together with reduced expression of markers of neurogenesis and synaptic development. Under baseline conditions, microRNA profiling and molecular analyses revealed increased miR-30c-5p expression and decreased Neurod1 mRNA and protein levels in the neonatal brain. RISC and luciferase reporter assays supported an interaction between miR-30c-5p and Neurod1 mRNA. In vivo administration of a miR-30c-5p mimic reduced Neurod1 expression and increased infarct size, whereas Neurod1 knockdown independently exacerbated ischemic injury and associated cellular changes. Conversely, inhibition of miR-30c-5p restored Neurod1 expression, reduced infarct size, and ameliorated associated pathological changes.

**Conclusions:** Maternal e-cigarette exposure is associated with baseline dysregulation of the miR-30c-5p–Neurod1 axis and increased susceptibility to neonatal ischemic brain injury in male rats. These findings suggest that this regulatory axis may contribute to the effects of prenatal e-cigarette exposure on the developing brain and may increase susceptibility to neonatal ischemic stroke.

## INTRODUCTION

Since their introduction into the US market approximately 2 decades ago, electronic cigarettes (e-cigarettes) have become increasingly popular, particularly among adolescents and women of reproductive age.^1–6^ Although commonly perceived as a safer alternative to conventional cigarettes and used for recreational purposes or smoking cessation, e-cigarette use has increased substantially among young women, including during pregnancy.^7–9^ A considerable proportion of pregnant women continue to use e-cigarette, raising concerns regarding their potential effects on fetal and offspring development.^7–9^ However, convincing evidence supporting the safety of e-cigarette use during pregnancy is lacking. Accumulating epidemiological and experimental evidence suggests that maternal e-cigarette exposure adversely affects fetal development and postnatal offspring health.^10–16^ Nevertheless, the effects of maternal e-cigarette exposure on neonatal brain vulnerability to ischemic injury and the underlying mechanisms remain poorly understood.

Neonatal arterial ischemic stroke (AIS) is a serious and often underrecognized neurological condition that can result in substantial long-term neurodevelopmental impairment.^17,18^ Unlike adult ischemic stroke, neonatal stroke exhibits distinct etiologies, risk factors, pathophysiological responses, and patterns of brain injury and recovery.^19–20^ Epidemiological and preclinical studies have identified maternal cigarette smoking and nicotine exposure as important risk factors associated with adverse fetal and neonatal outcomes, including hypoxic-ischemic encephalopathy (HIE) and ischemic brain injury.^21–24^ However, whether maternal e-cigarette exposure increases neonatal susceptibility to arterial ischemic brain injury and the mechanisms underlying this potential effect remain insufficiently characterized. Our previous studies demonstrated that maternal e-cigarette exposure exacerbates neonatal brain injury following both HIE and transient middle cerebral artery occlusion (tMCAO), a model of arterial ischemic brain injury.^25–26^ These findings suggest that prenatal e-cigarette exposure may reprogram neonatal brain vulnerability to ischemic insults. However, the molecular and epigenetic mechanisms underlying this heightened susceptibility remain largely unknown, limiting the development of targeted strategies to prevent or mitigate maternal e-cigarette exposure-associated neurological injury in offspring.

MicroRNAs (miRNAs) are small noncoding RNAs that regulate gene expression primarily through posttranscriptional mechanisms and represent an important component of epigenetic regulation.²⁷ In response to environmental and developmental stressors, altered miRNA expression can disrupt the regulation of target genes involved in fetal and neonatal brain development, potentially contributing to neurological dysfunction and disease susceptibility in offspring.^28–34^ Among these regulatory molecules, miR-30c-5p has emerged as an important modulator of cellular responses to ischemic injury, although its effects may vary depending on the cellular context and underlying pathological conditions.^35–38^ Neurod1 (neuronal differentiation 1), a transcription factor essential for neuronal differentiation, survival, and maturation, represents a potential downstream target relevant to these processes.^39^ Previous studies have demonstrated that Neurod1 deficiency increases susceptibility to ischemic brain injury, whereas Neurod1 restoration can ameliorate ischemic injury and neurodevelopmental deficits.^40,41^ Importantly, our previous mRNA sequencing analysis identified Neurod1 as a differentially expressed gene that was significantly downregulated in the neonatal brain following maternal e-cigarette exposure.^42^ However, whether miR-30c-5p regulates Neurod1 expression in the context of maternal e-cigarette exposure and whether this regulatory relationship contributes to neonatal ischemic brain injury remain unknown.

In the present study, we hypothesized that maternal e-cigarette exposure disrupts the miR-30c-5p–Neurod1 regulatory axis, thereby increasing neonatal susceptibility to ischemic brain injury and inducing additional pathological changes in the developing brain. To test this hypothesis, we investigated the effects of maternal e-cigarette exposure on neonatal brain injury following tMCAO and characterized associated changes in cellular senescence, apoptosis, neurogenesis-associated markers, and synaptic protein expression. We further examined the regulatory relationship between miR-30c-5p and Neurod1 using in vitro and in vivo approaches, assessed whether Neurod1 knockdown recapitulates the pathological effects of maternal e-cigarette exposure, and determined whether inhibition of miR-30c-5p ameliorates these adverse outcomes. These studies provide mechanistic insight into the role of the miR-30c-5p–Neurod1 axis in maternal e-cigarette exposure-associated neonatal brain vulnerability and identify a potential molecular pathway for future investigation of strategies to mitigate developmental neurological injury.

## METHODS

### Detailed experimental methods and procedures are provided in the Supplementary Materials

#### Data Availability Statement

The miRNA-seq data generated in this study are publicly available in the Gene Expression Omnibus under accession number GSE353596. The original contributions presented in the study are included within the article and the additional data supporting the findings of this study are available from the corresponding author upon reasonable request.

#### Animals and Ethical Approval

All animal procedures were approved by the Institutional Animal Care and Use Committee of Loma Linda University and conducted in accordance with the National Institutes of Health Guide for the Care and Use of Laboratory Animals. The study was conducted in accordance with the ARRIVE (Animal Research: Reporting of In Vivo Experiments) guidelines. Pregnant Sprague– Dawley rats were purchased from Charles River Laboratories (Portage, MI) and housed at a controlled temperature of 22°C under a 12-hour light/12-hour dark cycle, with food and water available ad libitum. Dams were randomly assigned to the e-cigarette exposure or control group. All experiments were conducted in neonatal offspring. Based on our previous findings that male offspring are more susceptible than female offspring to brain effects associated with perinatal nicotine and e-cigarette exposure,^23–25^ only male offspring were included in the present mechanistic studies.

#### Maternal E-cigarette Exposure Model

A chronic intermittent e-cigarette (CIEC) exposure model was used as we previously described.^25,26^ Pregnant Sprague–Dawley rats were randomly assigned to the e-cigarette exposure group (2.4% nicotine) or the control group and placed in comparable exposure chambers. E-cigarette aerosols were delivered using commercial BluPlus Cig e-cigarettes. Each exposure episode consisted of 6 puffs, with each puff lasting 4 seconds and a 30-second interval between puffs. One exposure episode was administered per hour during the 12-hour dark phase each day. This exposure regimen was designed to produce maternal blood nicotine and cotinine concentrations comparable to those reported in human e-cigarette users.^43–45^ Control dams underwent filtered-air exposure under the same chamber conditions and schedule as the e-cigarette group. Dams were exposed from embryonic day 4 (E4) through E20. After exposure, dams were returned to their home cages for spontaneous delivery. Offspring was used for experiments on postnatal day 10 (P10). At the designated experimental endpoint, animals were euthanized by inhalation of 5% isoflurane followed by brain or heart removal, as appropriate for the experiment. **Statistical Analysis**

Data are presented as mean ± SD. Statistical analyses were performed using GraphPad Prism 8 (GraphPad Software, San Diego, CA). Data normality was assessed using the Shapiro– Wilk test, and homogeneity of variance was evaluated using the F test. Comparisons between 2 groups were performed using an unpaired, 2-tailed Student’s t test for normally distributed data or the Mann–Whitney U test for nonnormally distributed data. Comparisons involving multiple groups or factors were performed using 1-way or 2-way analysis of variance (ANOVA), as appropriate, followed by Bonferroni post hoc multiple-comparison testing when indicated. A 2-tailed P value of less than 0.05 was considered statistically significant. The specific statistical tests used for each analysis are provided in the corresponding figure legends.

## RESULTS

### 1. Maternal e-cigarette exposure exacerbates tMCAO-induced ischemic brain injury and induces multiple pathological changes in the neonatal rat brain

Our previous studies demonstrated that prenatal nicotine/e-cigarette exposure sensitizes offspring to neonatal hypoxic-ischemic encephalopathy (HIE).^23,25,46,47^ Given the distinct pathological features and mechanisms underlying neonatal HIE and neonatal arterial ischemic brain injury, we further investigated whether maternal e-cigarette exposure increases susceptibility to transient middle cerebral artery occlusion (tMCAO)-induced ischemic brain injury in neonatal rats. Consistent with our previous findings, maternal e-cigarette exposure significantly increased infarct size following tMCAO, as determined by 2,3,5-triphenyltetrazolium chloride (TTC) staining (**Fig. 1A**). These findings indicate that prenatal e-cigarette exposure increases susceptibility to neonatal arterial ischemic brain injury.

**Figure 1.**
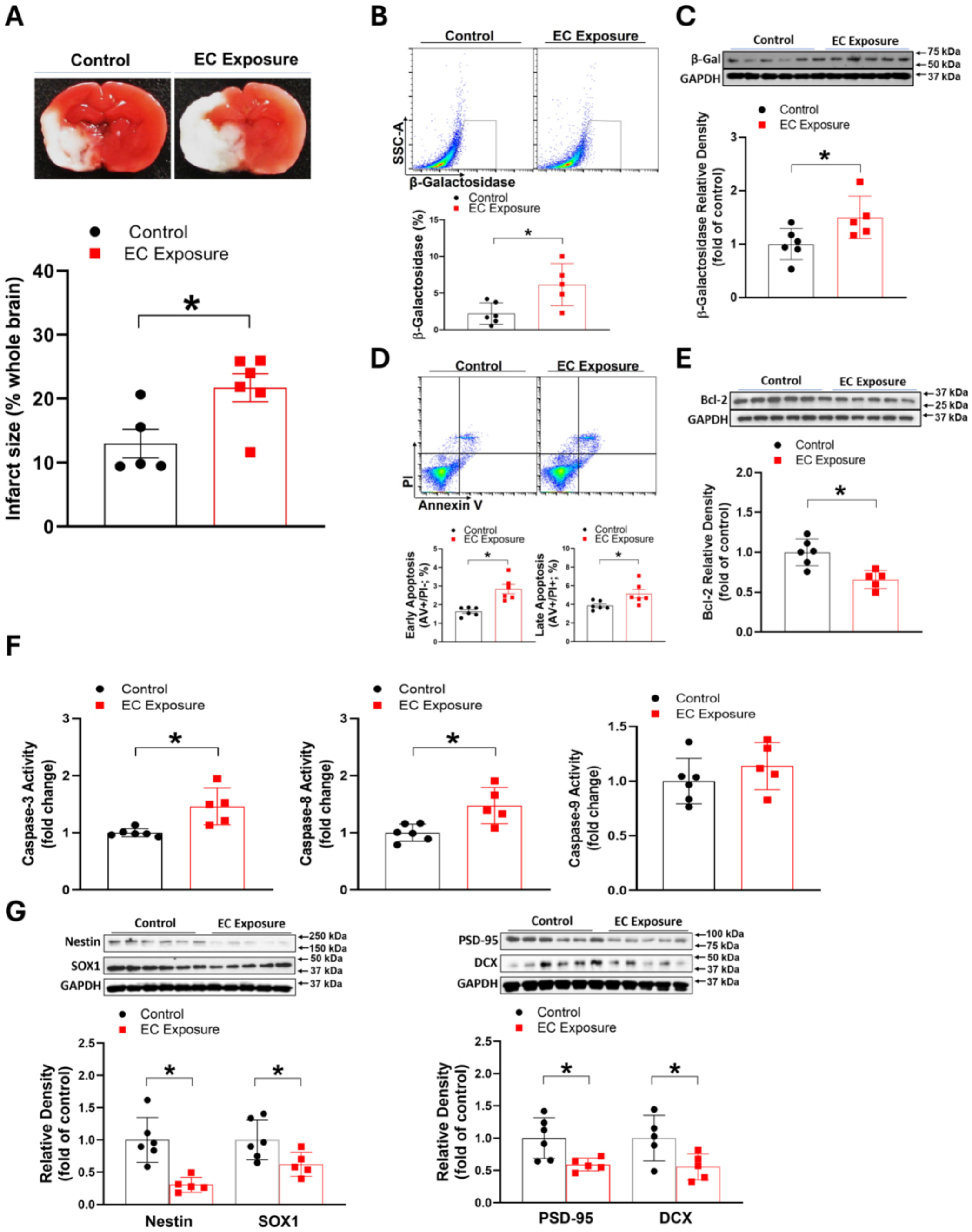
Maternal e-cigarette exposure exacerbates tMCAO-induced ischemic brain injury and induces multiple pathological changes in the neonatal rat brain. Rat pups were divided into 2 groups: control and maternal e-cigarette exposure. tMCAO was conducted on postnatal day 10 (P10), TTC staining was performed 48 hours after tMCAO. Brain tissues on P10 pups were collected for molecular studies. A, Quantification of tMCAO induced infarct size in neonatal rat brain via TTC staining. unpaired t-test, * *p<0.05,* (n=5-6). B, Flowcytometry quantification analysis of β-galactosidase levels in P10 neonatal brain. unpaired t-test, * *p<0.05,* (n=5-6). C, Representative blot images and quantification analysis of β-galactosidase protein abundance in P10 neonatal rat brain. unpaired t-test, * *p<0.05,* (n=5-6). D, Flowcytometry analysis of apoptosis levels in P10 neonatal brain. AV+/PI-indicates early apoptosis, AV+/PI+ indicates late apoptosis. unpaired t-test, * *p<0.05,* (n=6). E, Representative blot images and quantification analysis of anti-apoptotic Bcl-2 protein abundance in P10 neonatal rat brain. unpaired t-test, * *p<0.05,* (n=5-6). F, ELSIA quantification analysis of apoptosis activity of caspase 3, 8, 9, in P10 neonatal rat brain. unpaired t-test, * *p<0.05,* (n=5-6). G, Representative blot images and quantification analysis of Nestin, DCX, SOX1 and PSD-95 protein abundance in P10 neonatal rat brain. unpaired t-test, * *p<0.05,* (n=5-6). tMCAO, transit middle cerebral artery occlusion; TTC, 2,3,5-triphenyltetrazolium chloride; ELISA, enzyme-linked immunosorbent assay; DCX, doublecortin; SOX1, SRY-box transcription factor 1; PSD-95, postsynaptic density protein 95; AV, Annexin V; PI, propidium iodide.

We next examined additional pathological changes associated with maternal e-cigarette exposure. Flow cytometric analysis revealed an increased proportion of β-galactosidase-positive cells in the brains of offspring exposed to maternal e-cigarettes, accompanied by a significant increase in β-galactosidase protein abundance (**Fig. 1B** and **1C**), consistent with enhanced cellular senescence-associated changes. In addition, e-cigarette-exposed offspring exhibited an increased proportion of Annexin V-positive cells (**Fig. 1D**), reduced expression of the anti-apoptotic protein Bcl-2 (**Fig. 1E**), and increased caspase-3 and caspase-8 activities (**Fig. 1F**), indicating enhanced apoptotic responses. Western blot analysis further revealed significantly reduced expression of the neurogenesis-associated markers Nestin, DCX, and SOX1, as well as the synaptic structural protein PSD-95, in the neonatal brain following maternal e-cigarette exposure (**Fig. 1G**). Collectively, these findings demonstrate that maternal e-cigarette exposure exacerbates tMCAO-induced ischemic brain injury and is associated with multiple pathological alterations in the neonatal brain, including increased senescence-associated changes and apoptosis and reduced expression of neurogenesis-associated and synaptic proteins.

### 2. Maternal e-cigarette exposure increases miR-30c-5p expression and decreases Neurod1 expression in the neonatal rat brain

To investigate potential epigenetic mechanisms underlying the adverse effects of maternal e-cigarette exposure on the neonatal brain, we performed microRNA sequencing (miRNA-seq) using neonatal brain tissue. The sequencing analysis revealed differential expressions of miRNAs and piRNAs following maternal e-cigarette exposure, including increased expression of miR-30c-5p (**Fig. 2A**). Quantitative reverse-transcription polymerase chain reaction (qRT-PCR) confirmed the significant upregulation of miR-30c-5p in the brains of e-cigarette-exposed offspring (**Fig. 2B**).

**Figure 2.**
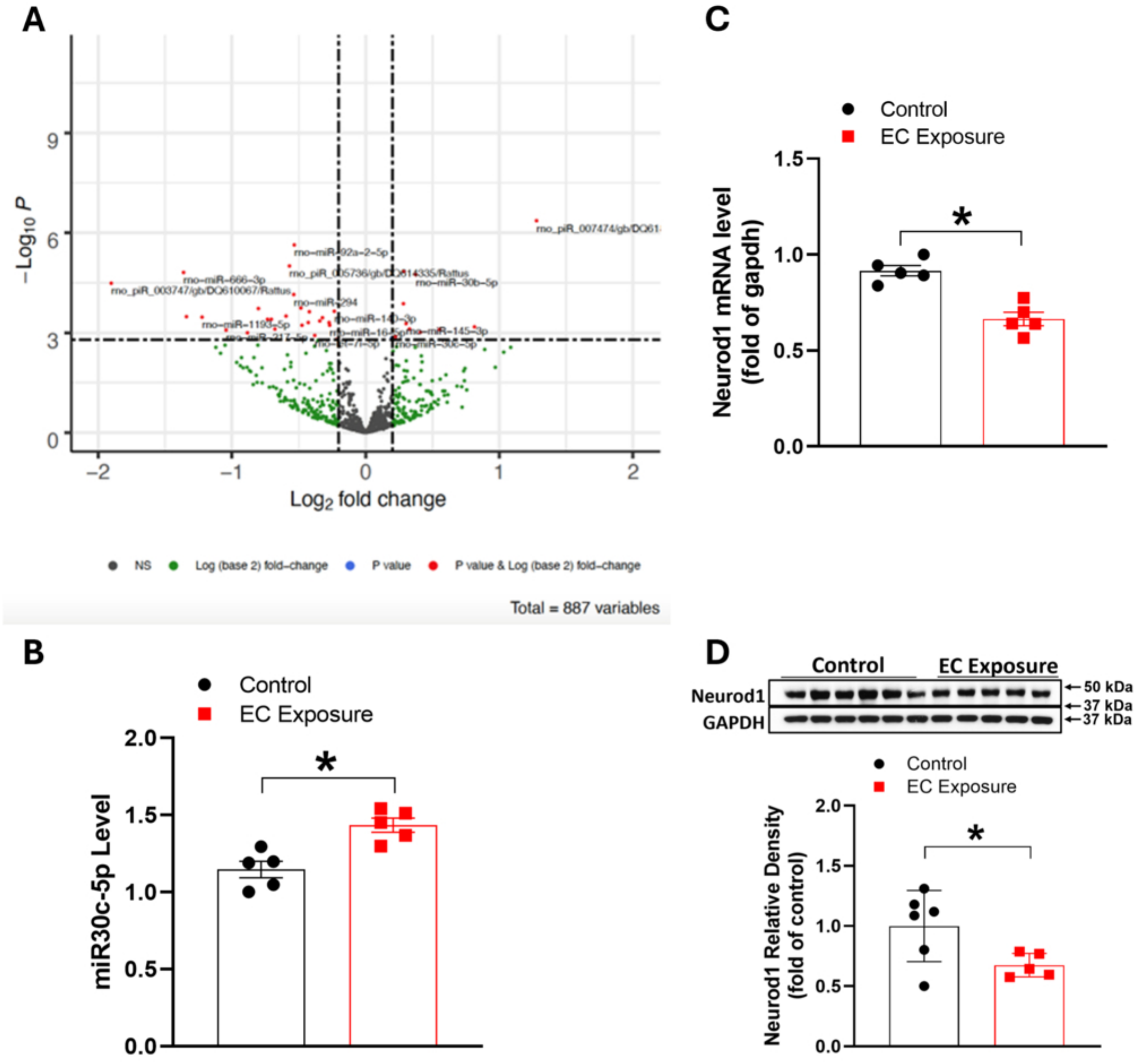
Maternal e-cigarette exposure increases miR-30c-5p expression and decreases Neurod1 expression in the neonatal rat brain. Rat pups were divided into 2 groups: control and maternal e-cigarette exposure. Brain tissues on P10 pups were collected for molecular studies. **A,** MiRNA-seq (volcano plot) reveals maternal e-cigarette exposure significantly upregulates miR-30c-5p levels in neonatal rat brain. (n=5). **B,** qRT-PCR quantification analysis of miR-30c-5p expression levels in P10 neonatal rat brain. unpaired t-test, * *p<0.05,* (n=5-6). **C,** qRT-PCR quantification analysis of Neurod1 mRNA expression levels in P10 neonatal rat brain. unpaired t-test, * *p<0.05,* (n=5). **D,** Representative blot images and quantification analysis of Neurod1 protein abundance in P10 neonatal rat brain. unpaired t-test, * *p<0.05,* (n=5-6). MiRNA-seq, miRNA sequencing; Neurod1, neuronal differentiation 1; qRT-PCR, quantitative reverse transcription polymerase chain reaction.

Our previous mRNA-seq analysis also revealed that maternal e-cigarette exposure significantly reduced Neurod1 mRNA expression in the neonatal brain.⁴² In the present study, qRT-PCR and Western blot analyses independently confirmed significantly decreased Neurod1 mRNA and protein levels, respectively, following maternal e-cigarette exposure (**Fig. 2C** and **2D**). Given the reported roles of miR-30c-5p and Neurod1 in brain development and neurological pathophysiology,³⁵˒³⁷⁻⁴¹ these findings suggest that dysregulation of miR-30c-5p and Neurod1 may contribute to the adverse effects of maternal e-cigarette exposure on neonatal brain vulnerability to ischemic injury.

### 3. MiR-30c-5p targets Neurod1 and increases tMCAO-induced infarct size

Given the concurrent upregulation of miR-30c-5p and downregulation of Neurod1 following maternal e-cigarette exposure, we investigated whether miR-30c-5p regulates Neurod1 expression and contributes to ischemic brain injury. To examine the interaction between miR-30c-5p and Neurod1, we performed RNA-induced silencing complex (RISC) immunoprecipitation in PC12 cells transfected with a miR-30c-5p mimic or a negative-control mimic. RISC immunoprecipitation analysis revealed significantly increased association of miR-30c-5p with Neurod1-associated RNA following miR-30c-5p mimic transfection (**Fig. 3A**). To test whether miR-30c-5p directly targets Neurod1 3’ UTR, we further conducted luciferase reporter assay in PC12 cells. We generated dual-luciferase reporter containing the wild-type (WT) or mutant (Mut) predicted binding sites. Rat PC12 cells were transfected with psiCHECK-2 Neurod1 3’ UTR or its mutant and treated with the miR-30c-5p mimic or the negative control. As shown in **Fig. 3B**, a significant decrease in luciferase activity was observed after co-transfection of miR-30c-5p mimic and psiCHECK-2 Neurod1 WT, but not in negative control or psiCHECK-2 Neurod1Mut, demonstrating a direct interaction of miR-30c-5p with the 3’ UTR of Neurod1 mRNA transcript. Consistent with above findings *in vitro*, qRT-PCR demonstrated significantly reduced Neurod1 mRNA expression in miR-30c-5p mimic-transfected cells (**Fig. 3C**), supporting a directly regulatory relationship between miR-30c-5p and Neurod1.

**Figure 3.**
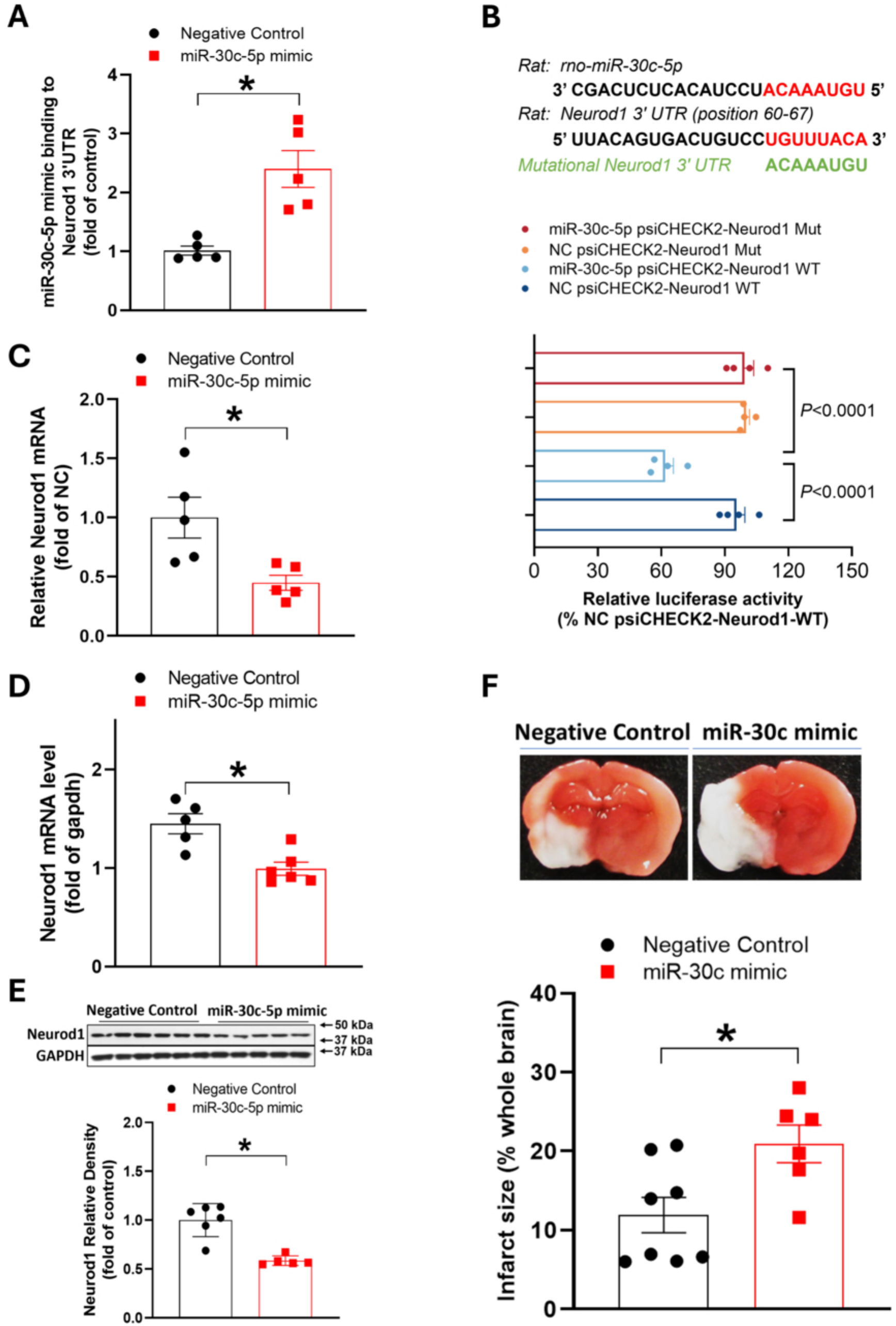
MiR-30c-5p directly targets Neurod1 and increases tMCAO-induced infarct size. *In vitro*, RISC-IP and luciferase reporter assay were conducted to determine whether miR-30c-5p directly targets Neurod1. PC-12 cells were transfected with miR-30c-5p mimic or negative control in RISC-IP. In luciferase reporter assay, PC-12 cells were transfected with psiCHECK-2 Neurod1 3’ UTR or psiCHECK-2 Neurod1 mutant together with miR-30c-5p mimic or negative control. *In vivo*, miR-30c-5p mimic or negative control ICV was administered on P8 naïve pups, tMCAO conducted on P10 and brain tissues on P10 were collected for molecular studies. **A,** RISC-IP revealed there was significantly increased binding capacity of miR-30c-5p mimic to Neurod1 3’ UTR. unpaired t-test, * *p<0.05,* (n=5 independent experiment). **B,** Luciferase reporter assay demonstrated there was a significant decrease in luciferase activity after co-transfection of miR-30c-5p mimic and psiCHECK-2 Neurod1 WT. unpaired t-test (n=4 independent experiment). **C,** qRT-PCR quantification analysis of Neurod1 mRNA expression levels in PC-12 cells. unpaired t-test, * *p<0.05,* (n=5 independent experiment). **D,** qRT-PCR quantification analysis of Neurod1 mRNA expression levels in P10 pup brain subjected to miR-30c-5p mimic or negative control ICV. unpaired t-test, * *p<0.05,* (n=5-6 pups). **E,** Representative blot images and quantification analysis of Neurod1 protein abundance in P10 neonatal rat brain. unpaired t-test, * *p<0.05,* (n=5-6). **F,** Quantification of tMCAO induced infarct size in neonatal rat brain subjected to miR-30c-5p mimic or negative control ICV via TTC staining. unpaired t-test, * *p<0.05,* (n=6-8 pups). RISC-IP, RNA-induced silencing complex-immunoprecipitation; ICV, intracerebroventricular.

We next examined this relationship *in vivo* by administering a miR-30c-5p mimic intracerebroventricularly (ICV) to naïve postnatal day 8 (P8) rat pups. Consistent with the *in vitro* findings, miR-30c-5p mimic administration significantly reduced Neurod1 mRNA and protein levels in the neonatal brain (**Fig. 3D** and **3E**). Importantly, miR-30c-5p mimic pretreatment also significantly increased tMCAO-induced infarct size (**Fig. 3F**). Together, these findings support a functional miR-30c-5p–Neurod1 regulatory axis and suggest that increased miR-30c-5p expression contributes to heightened susceptibility to ischemic brain injury in neonatal rats.

### 4. Neurod1 knockdown recapitulates pathological changes associated with maternal e-cigarette exposure in the neonatal rat brain

Neurod1 is an important regulator of brain development and neuronal maturation.³⁹⁻⁴¹ Given our finding that maternal e-cigarette exposure significantly reduces Neurod1 expression in the neonatal brain, we investigated whether Neurod1 downregulation contributes to the associated pathological changes. We used Neurod1-targeting small interfering RNA (siRNA) to reduce Neurod1 expression in naïve neonatal rats and assessed the resulting effects on ischemic injury and brain pathology.

Neurod1 knockdown significantly reduced Neurod1 protein expression and increased tMCAO-induced infarct size in neonatal rats (**Fig. 4A**). Furthermore, Neurod1 knockdown increased the proportion of β-galactosidase-positive cells, as determined by flow cytometry (**Fig. 4B**), and increased β-galactosidase protein abundance (**Fig. 4C**), consistent with enhanced senescence-associated changes. Flow cytometric analysis also revealed increased proportions of cells in both the early apoptotic (Annexin V-positive/propidium iodide-negative [AV⁺/PI⁻]) and late apoptotic (AV⁺/PI⁺) populations (**Fig. 4D**). These changes were accompanied by reduced Bcl-2 protein expression (**Fig. 4E**). In addition, Neurod1 knockdown significantly decreased the expression of the neurogenesis-associated markers Nestin, DCX, and SOX1, as well as the synaptic structural protein PSD-95 (**Fig. 4F**). These findings indicate that Neurod1 downregulation recapitulates several pathological changes associated with maternal e-cigarette exposure, including increased susceptibility to ischemic injury, enhanced senescence-associated and apoptotic changes, and reduced expression of neurogenesis-associated and synaptic proteins. Collectively, these results support a protective role for Neurod1 in maintaining neonatal brain homeostasis.

**Figure 4.**
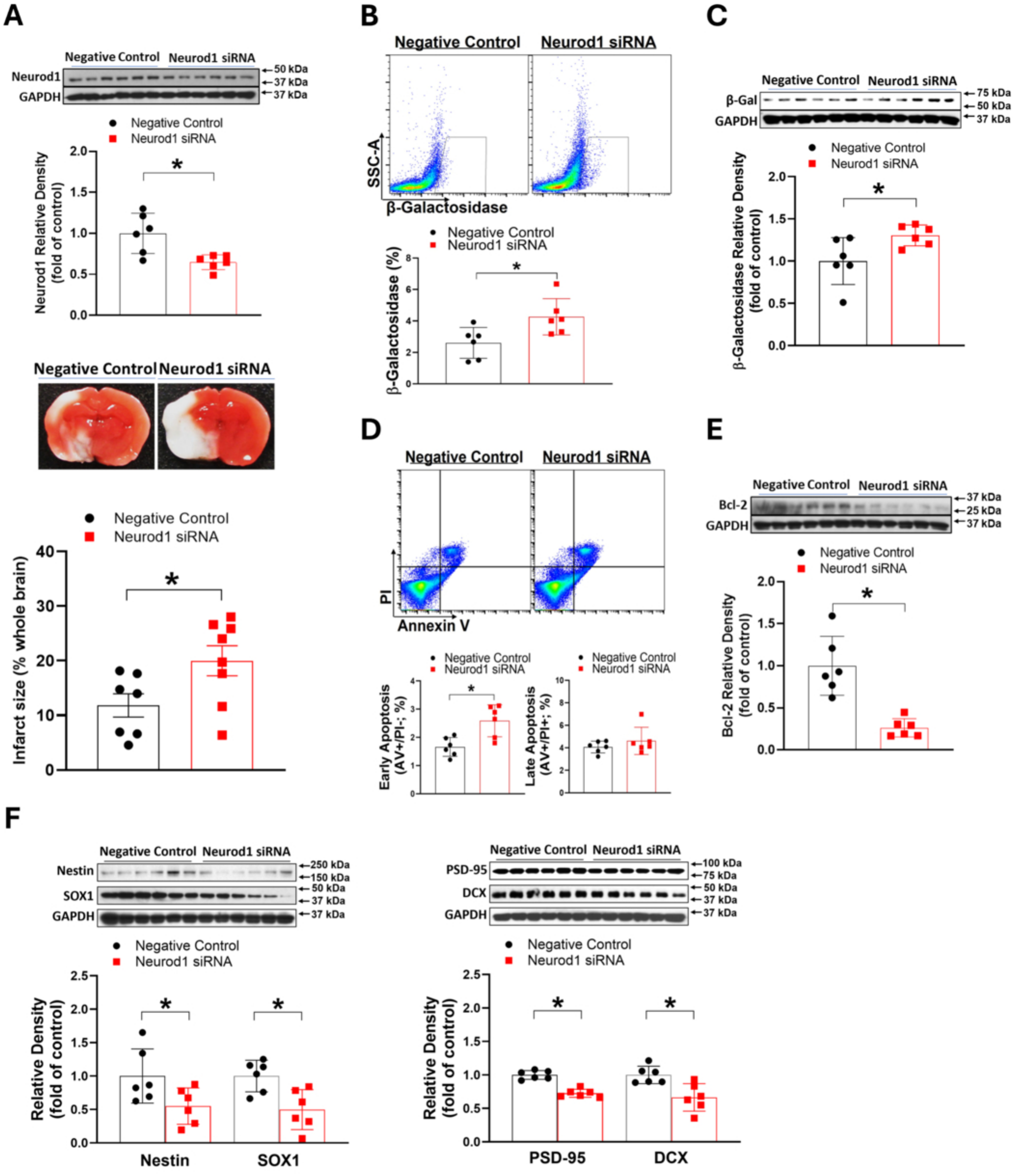
Neurod1 knockdown recapitulates pathological changes associated with maternal e-cigarette exposure in the neonatal rat brain. Naïve neonatal rat pups were divided into 2 groups subjected to ICV Neurod1 siRNA or its negative control on P8, respectively. tMCAO was conducted on P10. TTC staining was performed on P12. Brain tissues on P10 were collected for molecular studies. **A,** Representative blot images and quantification analysis of Neurod1 protein abundance in P10 neonatal rat brain. unpaired t-test, * *p<0.05,* (n=6). And quantification of tMCAO induced infarct size in neonatal rat brain subjected to ICV Neurod1 siRNA or its negative control via TTC staining. unpaired t-test, * *p<0.05,* (n=7-8). **B,** Flowcytometry quantification analysis of β-galactosidase levels in P10 neonatal brain. unpaired t-test, * *p<0.05,* (n=6). **C,** Representative blot images and quantification analysis of β-galactosidase protein abundance in P10 neonatal rat brain. unpaired t-test, * *p<0.05,* (n=6). **D,** Flowcytometry analysis of apoptosis levels in P10 neonatal brain. AV+/PI-indicates early apoptosis, AV+/PI+ indicates late apoptosis. unpaired t-test, * *p<0.05,* (n=6). **E,** Representative blot images and quantification analysis of anti-apoptotic Bcl-2 protein abundance in P10 neonatal rat brain. unpaired t-test, * *p<0.05,* (n=6). **F,** Representative blot images and quantification analysis of Nestin, DCX, SOX1 and PSD-95 protein abundance in P10 neonatal rat brain. unpaired t-test, * *p<0.05,* (n=6).

### 5. Inhibition of miR-30c-5p normalizes Neurod1 expression and attenuates maternal e-cigarette exposure-associated ischemic brain injury and pathological changes

Our findings indicate that Neurod1 knockdown recapitulates multiple pathological changes associated with maternal e-cigarette exposure and that miR-30c-5p negatively regulates Neurod1 expression. We therefore investigated whether inhibiting miR-30c-5p could restore Neurod1 expression and mitigate the adverse effects of maternal e-cigarette exposure on the neonatal brain. To this end, we administered a locked nucleic acid (LNA) inhibitor targeting miR-30c-5p intracerebroventricularly.

As shown in **Fig. 5**, inhibition of miR-30c-5p abolished the maternal e-cigarette exposure-associated reductions in Neurod1 mRNA and protein expression (**Fig. 5A** and **5B**). Importantly, miR-30c-5p inhibition also normalized the increased infarct size following transient middle cerebral artery occlusion (tMCAO) in offspring exposed to maternal e-cigarettes (**Figure 5C**).

**Figure 5.**
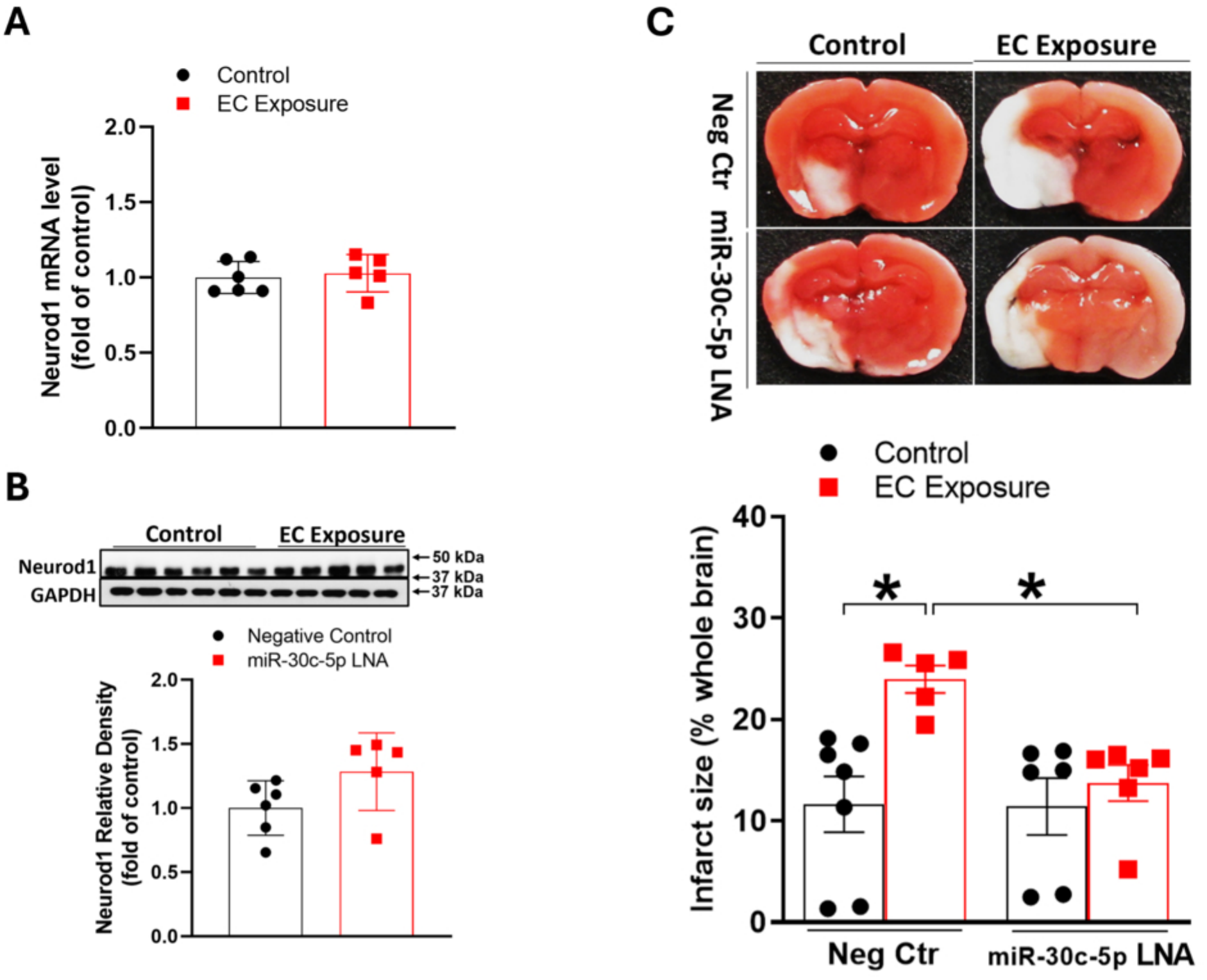
Inhibition of miR-30c-5p restores Neurod1 level and attenuates maternal e-cigarette exposure induced exacerbation of ischemic brain injury in neonatal rat brain. Neonatal rat pups were divided into 2 groups: control and maternal e-cigarette exposure. miR-30c-5p LNA was administered via ICV on P8 for miR-30c-5p inhibition. tMCAO was conducted on P10. TTC staining on P12. Brain tissues on P10 pups were collected for molecular studies. **A,** qRT-PCR quantification analysis of Neurod1 mRNA expression levels in P10 pup brain subjected to miR-30c-5p LNA ICV. unpaired t-test (n=6). **B,** Representative blot images and quantification analysis of Neurod1 protein abundance in P10 neonatal rat brain subjected to miR-30c-5p LNA ICV. unpaired t-test (n=6). **C,** Quantification of tMCAO induced infarct size in neonatal rat brain subjected to miR-30c-5p LNA ICV via TTC staining. unpaired t-test, * *p<0.05,* (n=5-7 pups). LNA, locked nucleic acid.

We next investigated whether miR-30c-5p inhibition could reverse the additional pathological changes associated with maternal e-cigarette exposure. As shown in **Figure 6**, miR-30c-5p inhibition abolished the increases in senescence-associated measures, including the proportion of β-galactosidase-positive cells and β-galactosidase protein abundance (**Fig. 6A** and **6B**). It also normalized apoptosis-associated changes, as assessed by apoptotic cell populations and apoptosis-related protein measures (**Fig. 6C–6E**). Furthermore, miR-30c-5p inhibition reversed the e-cigarette exposure-associated alterations in the expression of neurogenesis-associated markers and the synaptic protein PSD-95 (**Fig. 6F**).

**Figure 6.**
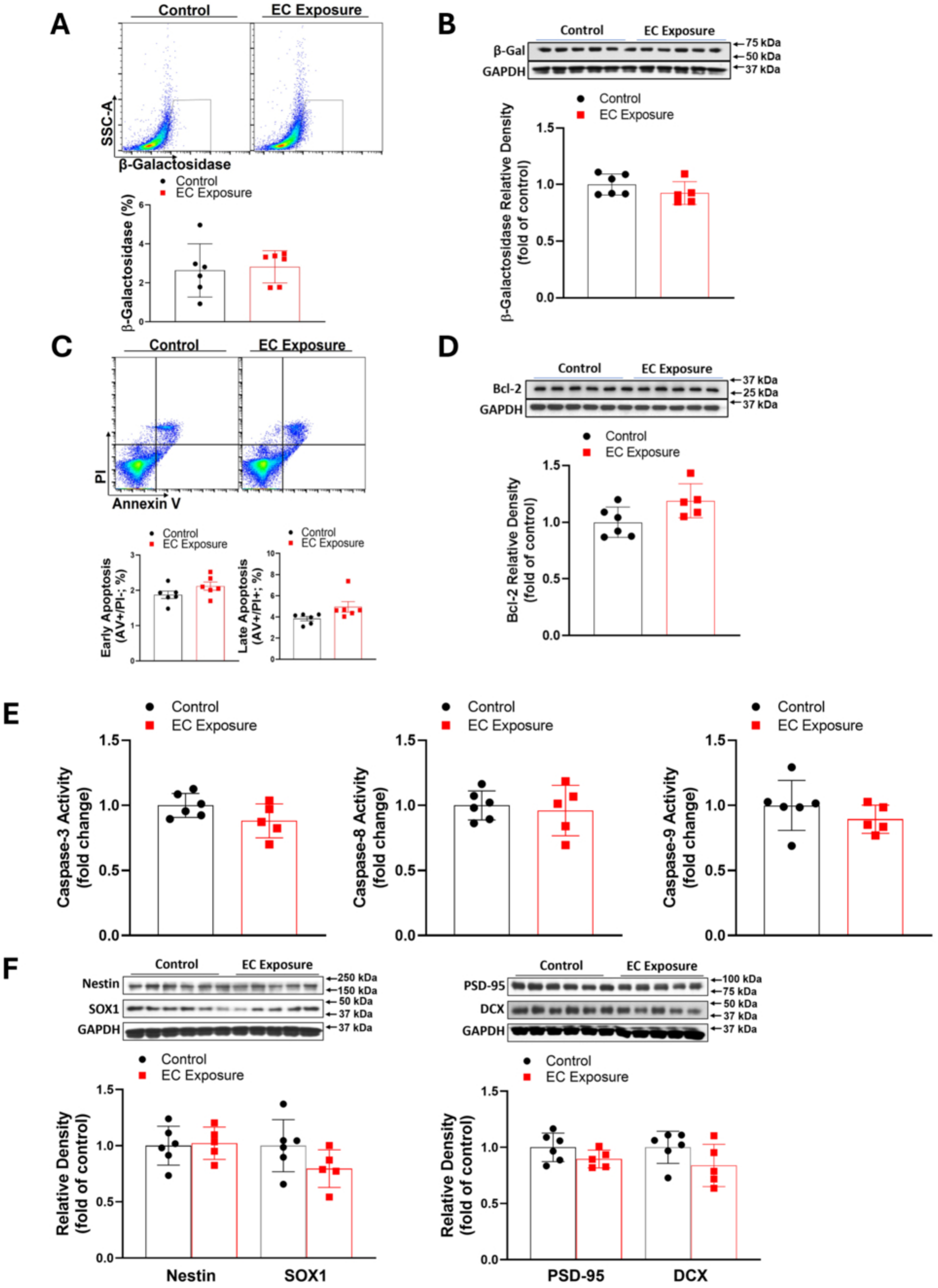
Inhibition of miR-30c-5p abrogates maternal e-cigarette exposure induced multiple pathological changes in neonatal rat brain. Neonatal rat pups were divided into 2 groups: control and maternal e-cigarette exposure. miR-30c-5p LNA was administered via ICV on P8 for miR-30c-5p inhibition. Brain tissues on P10 pups were collected for molecular studies. **A,** Flowcytometry quantification analysis of β-galactosidase levels in P10 neonatal brain subjected to miR-30c-5p LNA ICV. unpaired t-test (n=6). **B,** Representative blot images and quantification analysis of β-galactosidase protein abundance in P10 neonatal rat brain subjected to miR-30c-5p LNA ICV. unpaired t-test (n=5-6). **C,** Flowcytometry analysis of apoptosis levels in P10 neonatal brain subjected to miR-30c-5p LNA ICV. AV+/PI-indicates early apoptosis, AV+/PI+ indicates late apoptosis. unpaired t-test (n=6). **D,** Representative blot images and quantification analysis of anti-apoptotic Bcl-2 protein abundance in P10 neonatal rat brain subjected to miR-30c-5p LNA ICV. unpaired t-test (n=5-6). **E,** ELSIA quantification analysis of apoptosis activity of caspase 3, 8, 9, in P10 neonatal rat brain subjected to miR-30c-5p LNA ICV. unpaired t-test (n=5). **F,** Representative blot images and quantification analysis of Nestin, DCX, SOX1 and PSD-95 protein abundance in P10 neonatal rat brain subjected to miR-30c-5p LNA ICV. unpaired t-test (n=6).

Collectively, these findings demonstrate that inhibition of miR-30c-5p reverses multiple adverse effects associated with maternal e-cigarette exposure, including reduced Neurod1 expression, increased susceptibility to ischemic brain injury, and pathological alterations involving senescence-associated changes, apoptosis, neurogenesis-associated markers, and synaptic protein expression. These results support a functional role for the miR-30c-5p–Neurod1 regulatory axis in mediating the adverse effects of maternal e-cigarette exposure on neonatal brain vulnerability to ischemic injury.

## DISCUSSION

The present study provides preclinical evidence that maternal e-cigarette exposure exacerbates transient middle cerebral artery occlusion (tMCAO)-induced ischemic brain injury and is associated with multiple pathological alterations in the neonatal brain. The major findings are as follows: (1) maternal e-cigarette exposure increased infarct size following tMCAO; (2) maternal e-cigarette exposure was associated with increased apoptosis and senescence-associated changes and reduced expression of neurogenesis-associated markers and synaptic proteins; (3) miR-30c-5p was upregulated, whereas Neurod1 expression was reduced, in the neonatal brain following maternal e-cigarette exposure; (4) experimental manipulation of miR-30c-5p and Neurod1 supported a functional role for this regulatory pathway in ischemic injury and associated brain pathology; and (5) inhibition of miR-30c-5p restored Neurod1 expression and attenuated multiple adverse outcomes associated with maternal e-cigarette exposure. Together, these findings identify the miR-30c-5p–Neurod1 axis as a potential molecular pathway linking prenatal e-cigarette exposure to increased neonatal brain vulnerability to ischemic injury.

Been marketed as a safer alternative to conventional cigarettes, e-cigarettes have gained highly popularity in the United States. ^1–9^ However, no convincing evidence suggests e-cigarettes are safe to be used, especially during pregnancy. Given that e-cigarette aerosols contain nicotine and other potentially harmful constituents, including propylene glycol, vegetable glycerin, metals, and flavoring-related chemicals, emerging evidence reveals that maternal e-cigarette exposure can adversely affect fetal and offspring development.^10–16^ Our previous studies demonstrated that prenatal nicotine exposure reprograms brain development through epigenetic mechanisms, including alterations in DNA methylation and miRNA expression.^23,24,46,47^ More recent studies from our group showed that maternal e-cigarette exposure alters DNA methylation and transcriptomic profiles, disrupts metabolic and developmental programs in the neonatal brain, and exacerbates neonatal hypoxic-ischemic encephalopathy (HIE).^25,42,48^ However, whether maternal e-cigarette exposure also increases susceptibility to arterial ischemic brain injury and the underlying molecular mechanisms have remained insufficiently characterized. Because HIE and arterial ischemic stroke involve distinct pathophysiological processes, ^49,50^ we investigated the effects of maternal e-cigarette exposure using a neonatal tMCAO model. We found that maternal e-cigarettes exposure significantly increased infarct size and was accompanied by multiple pathological alterations in the neonatal brain. These findings extend our previous observations beyond HIE and suggest that prenatal e-cigarette exposure may increase neonatal susceptibility to different forms of ischemic brain injury. Furthermore, our mechanistic experiments identify the miR-30c-5p–Neurod1 regulatory axis as a potential contributor to these adverse effects.

Neurod1 is a transcription factor with essential roles in neuronal differentiation, maturation, and survival.³⁹˒⁴⁰ It also participates in neuronal reprogramming and has attracted interest as a potential target for regenerative approaches to neurological injury.^41,51–53^ Preclinical studies have reported that Neurod1-based strategies can promote neuronal regeneration and improve outcomes in experimental models of ischemic stroke, spinal cord injury, traumatic brain injury, and neurodegenerative disease.^41,54–58^ Consistent with these findings, our previous transcriptomic analysis identified Neurod1 as a gene downregulated in the neonatal brain following maternal e-cigarette exposure.⁴² In the present study, we confirmed reduced Neurod1 mRNA and protein expression and demonstrated that Neurod1 knockdown increased tMCAO-induced infarct size. Neurod1 knockdown also reproduced several pathological changes associated with maternal e-cigarette exposure, including increased senescence-associated measures and apoptotic responses and reduced expression of neurogenesis-associated markers and PSD-95. These findings support an important role for Neurod1 in maintaining neonatal brain homeostasis and suggest that reduced Neurod1 expression may contribute to the adverse effects of prenatal e-cigarette exposure. Nevertheless, because Neurod1 knockdown does not reproduce every possible consequence of maternal exposure, other molecular pathways are likely to contribute to the observed phenotype. Further studies are needed to determine the cell-specific functions of Neurod1 and establish how its downregulation affects neuronal development and the response to ischemic injury.

An important finding of this study is the identification of miR-30c-5p as a potential upstream regulator of Neurod1 in the context of maternal e-cigarette exposure. miRNAs regulate gene expression post-transcriptionally and are responsive to developmental and environmental stressors, making them plausible mediators of developmental programming.²⁷⁻³⁴ Previous studies have implicated miR-30c-5p in ischemic and other pathological processes, although its reported effects vary according to the experimental model and biological context.³⁵⁻³⁸ In our study, miRNA sequencing and qRT-PCR demonstrated increased miR-30c-5p expression in the neonatal brain following maternal e-cigarette exposure, whereas Neurod1 expression was reduced. Both RISC immunoprecipitation and luciferase reporter assay in PC12 cells supported the direct interaction between miR-30c-5p and Neurod1-associated RNA, and experimental administration of a miR-30c-5p mimic reduced Neurod1 expression in vitro and in vivo. Moreover, miR-30c-5p mimic administration increased tMCAO-induced infarct size. These findings support a functional regulatory relationship between miR-30c-5p and Neurod1 and suggest that increased miR-30c-5p expression may contribute to neonatal ischemic vulnerability. Importantly, inhibition of miR-30c-5p restored Neurod1 expression and attenuated the increased infarct size and other pathological changes associated with maternal e-cigarette exposure. The complementary gain- and loss-of-function findings strengthen the evidence for a role of miR-30c-5p in this process. However, because miRNAs can regulate multiple target genes, additional experiments are needed to determine the extent to which the protective effects of miR-30c-5p inhibition are specifically mediated by Neurod1.

The findings also highlight the importance of developmental context in interpreting the role of miR-30c-5p in ischemic brain injury. Some previous studies have reported neuroprotective effects of miR-30c-5p in experimental ischemia.³⁷˒³⁸ At first glance, these observations appear to contrast with our finding that increased miR-30c-5p expression is associated with greater ischemic injury in neonatal rats. Several factors may account for this difference. First, neonatal and adult brains differ in cellular composition, developmental programs, and responses to ischemia.¹⁸⁻²⁰ Second, prenatal e-cigarette exposure may alter the molecular and epigenetic environment of the developing brain before an ischemic insult occurs. Thus, the consequences of miR-30c-5p dysregulation may depend on developmental stage, pre-existing molecular changes, and the availability of relevant target genes. Our findings therefore suggest that the function of miR-30c-5p in ischemic injury should be interpreted within the specific developmental and exposure context rather than assumed to be uniformly protective or detrimental. Further comparative studies in neonatal and adult models will be necessary to determine the mechanisms underlying these context-dependent effects.

In addition to increased ischemic injury, maternal e-cigarette exposure was associated with several pathological changes in the neonatal brain, including increased apoptosis, senescence-associated changes, and reduced expression of neurogenesis-associated markers and PSD-95. These findings suggest that prenatal exposure may affect the developing brain beyond its response to an acute ischemic insult. Reduced expression of Nestin, DCX, and SOX1 is consistent with altered neurogenesis-associated processes, whereas decreased PSD-95 suggests potential disruption of synaptic development or maintenance. Increased β-galactosidase-positive cell proportions and protein abundance are consistent with senescence-associated changes, although these measures alone do not establish the cellular identity or functional consequences of senescence. Similarly, the apoptosis-related findings indicate enhanced apoptotic responses but do not fully define the affected cell populations. Notably, Neurod1 knockdown reproduced several of these changes, and miR-30c-5p inhibition attenuated them in e-cigarette-exposed offspring. This convergence supports the involvement of the miR-30c-5p–Neurod1 pathway in multiple aspects of neonatal brain pathology. Because these outcomes were assessed primarily through molecular and cellular measures, further studies incorporating cell-specific analyses, functional assessments, and long-term behavioral outcomes are needed to determine whether the observed changes translate into persistent neurodevelopmental deficits or increased susceptibility to neurological disease later in life.

The present findings have potential translational implications. Neonatal arterial ischemic stroke can result in substantial long-term neurological morbidity, yet its prenatal determinants and molecular mechanisms remain incompletely understood.¹⁸⁻²⁰ Our results suggest that maternal e-cigarette exposure may be an important modifiable exposure to investigate in relation to neonatal ischemic vulnerability. The miR-30c-5p–Neurod1 axis may also provide a starting point for developing molecular approaches to understand or mitigate exposure-associated brain injury. However, the current study establishes preclinical evidence in a rat model and does not demonstrate that miR-30c-5p inhibition is safe or effective in human neonates. The translational relevance of this pathway will require validation in additional experimental models and human studies, together with careful evaluation of developmental safety, treatment timing, and potential off-target effects. These findings reinforce the importance of avoiding e-cigarette exposure during pregnancy while highlighting the need for further research into its effects on neonatal brain development and ischemic injury.

Several limitations should be acknowledged. First, the present experiments were conducted exclusively in male neonatal rats, limiting the generalizability of the findings to female offspring. Our previous studies suggest that female offspring may exhibit different responses to prenatal nicotine or e-cigarette exposure, highlighting the need to investigate potential sex-dependent mechanisms. Second, most mechanistic analyses were performed using whole-brain tissue. Given the cellular and regional heterogeneity of the brain, further studies should determine whether the miR-30c-5p–Neurod1 pathway is altered in specific neuronal or non-neuronal populations and identify the brain regions most affected by maternal exposure. Third, although the use of miRNA mimics, LNA inhibitors, and siRNA knockdown provided complementary evidence for the involvement of this pathway, these approaches have limitations related to target specificity, delivery, and potential off-target effects. Future studies using more selective genetic approaches, including targeted Neurod1 manipulation and cell-specific interventions, will help clarify the causal relationship between miR-30c-5p, Neurod1, and the observed pathological outcomes. In particular, restoring Neurod1 expression independently of miR-30c-5p inhibition would help determine whether Neurod1 is sufficient to mediate the protective effects of pathway manipulation. Fourth, the study focused primarily on acute ischemic injury and early pathological changes. Longitudinal studies are needed to determine whether maternal e-cigarette exposure produces persistent neurological deficits and whether intervention targeting this pathway improves long-term structural and functional outcomes. Finally, although our findings implicate the miR-30c-5p– Neurod1 axis, the molecular mechanisms linking maternal e-cigarette exposure to miR-30c-5p upregulation remain unresolved and warrant further investigation.

In summary, this study provides preclinical evidence that maternal e-cigarette exposure exacerbates tMCAO-induced ischemic brain injury and is associated with multiple pathological changes in the neonatal rat brain. Our findings support a functional role for the miR-30c-5p– Neurod1 regulatory axis in these effects, as Neurod1 knockdown reproduced several exposure-associated phenotypes, whereas miR-30c-5p inhibition restored Neurod1 expression and attenuated ischemic injury and associated pathological changes. These results identify the miR-30c-5p–Neurod1 axis as a potential mechanism linking prenatal e-cigarette exposure to neonatal brain vulnerability. Further studies are needed to establish its cell-specific functions, determine its contribution to long-term neurological outcomes, and assess its translational potential.

## Acknowledgments

We would like to thank Maryam Hosseini Asanjan for her assistance with preparation of miRNA sequencing data.

## Supplemental Material

Supplemental Methods

ARRIVE Checklist

## Non-standard Abbreviations and Acronyms

AIS: arterial ischemic stroke
AV: Annexin V
CIEC: chronic intermittent e-cigarette
DCX: doublecortin
DEGs: differentially expressed genes
ELISA: enzyme-linked immunosorbent assay
HIE: hypoxic ischemic encephalopathy
ICV: intracerebroventricular
LNA: locked nucleic acid
Neurod1: neuronal differentiation 1
PI: propidium iodide
PSD-95: postsynaptic density protein 95
qRT-PCR: quantitative reverse transcription polymerase chain reaction
RISC: RNA-induced silencing complex
siRNA: small interfering
RNA SOX1: SRY-box transcription factor 1
tMCAO: transient middle cerebral artery occlusion
TTC: 2,3,5-triphenyltetrazolium chloride

